# Extramacrochaete promotes branch and bouton number via the sequestration of Daughterless in the cytoplasm of neurons

**DOI:** 10.1101/686014

**Authors:** Edward A. Waddell, Jennifer M. Viveiros, Erin L. Robinson, Michal A. Sharoni, Nina K. Latcheva, Daniel R. Marenda

## Abstract

The class I basic Helix Loop Helix (bHLH) proteins are highly conserved transcription factors that are ubiquitously expressed. A wealth of literature on class I bHLH proteins have shown that these proteins must homodimerize or heterodimerize with tissue specific HLH proteins in order to bind DNA at E box (CANNTG) consensus sequences to control tissue specific transcription. Due to its ubiquitous expression, class I bHLH proteins are also extensively regulated post-translationally, mostly through dimerization. Previously, we reported that in addition to its role in promoting neurogenesis, the class I bHLH protein Daughterless also functions in mature neurons to restrict axon branching and synapse number. Here, we show that part of the molecular logic that specifies how Daughterless functions in neurogenesis is also conserved in neurons. We show that the type V HLH protein Extramacrochaete binds to and represses Daughterless function by sequestering Daughterless to the cytoplasm. This work provides initial insights into the mechanisms underlying the function of Daughterless and Extramacrochatae in neurons while providing a novel understanding of how Extramacrochaetae functions to restricts Daughterless activity within the cell.

## Introduction

The class I basic Helix Loop Helix (bHLH) proteins are transcription factors that are highly conserved from invertebrates to humans, expressed in a broad number of tissues during development, and generally recognized as master regulators of neurogenesis (reviewed in (Castro and Guillemot, 2011; Guillemot, 2007; Powell and Jarman, 2008)). Extensive literature on class I bHLH proteins have shown that these proteins are ubiquitously expressed, and must homodimerize or heterodimerize with tissue specific HLH proteins in order to bind DNA at E box (CANNTG) consensus sequences to control tissue specific transcription (Cabrera and Alonso, 1991; Cronmiller and Cummings, 1993; Massari and Murre, 2000; Murre et al., 1989a; Smith and Cronmiller, 2001). This homo- or heterodimerization is critical for all class I bHLH proteins studied to date (Murre et al., 1989a; Murre et al., 1989b). Thus, much research has gone in to determining which binding partners help control the specific cellular functions of the type I bHLH proteins, as these partners are not only critical for determining the specificity of type I bHLH function, but also make attractive therapeutic targets for the multiple diseases that the class I bHLH proteins are associated with.

In *Drosophila*, Daughterless (Da) is the only class I bHLH protein present (Cronmiller and Cummings, 1993; Cronmiller et al., 1988). Until recently, Da function in the nervous system was thought to be restricted to controlling neurogenesis. However, we have recently shown that Da is also present in postmitotic cells (neurons) where it restricts synaptic branching and synapse number (D’Rozario et al., 2016). We have also demonstrated that the Da mammalian homolog, Transcription Factor 4 (Tcf4), functions similarly (D’Rozario et al., 2016). In order to mediate this synaptic restriction, Da and Tcf4 bind to cis regulatory regions of the *neurexin* gene and repress *neurexin* transcription in neurons and not in proliferating precursor cells.

Here, we report that part of the molecular logic that specifies how Da functions in mitotically active cells is conserved in neurons via the binding and subsequent repression of Da function by the type V HLH protein Extramacrochaete (Emc). In mitotic cells, Emc serves as a negative feedback regulator of *da* expression and Da activity (Campuzano, 2001; Garrell and Modolell, 1990; Spratford and Kumar, 2015b; van Doren et al., 1991). Emc lacks a basic domain, allowing it to heterodimerize with Da, but prevents Da from binding DNA and dimerizing to other transcription factors (Cabrera et al., 1994; Ellis et al., 1990). Recently, Spratford and Kumar have shown that Emc does this by sequestering Da away from DNA, thus preventing Da-mediated transcription (Spratford and Kumar, 2015b). Here, we expand upon this observation, and show that Emc accomplishes this sequestration predominantly by sequestering Da within the cytoplasm of neurons. We have determined that Emc is present in differentiated motor neurons of the *Drosophila* neuromuscular junction (NMJ), where it functions to promote axonal arborization and synapse number in these neurons. Emc accomplishes this by binding and inhibiting Da from repressing *neurexin* expression. Taken together, these data lend insight into the molecular functions of bHLH proteins in mature neurons and begins to tease apart the molecular logical that bHLH proteins use to control novel functions in postmitotic cells. Further, these data may help shed light on diseases associated with HLH proteins in humans.

## RESULTS

### Emc is present in neurons and promotes axonal branching and bouton number

In order to begin our analysis, we first wanted to determine whether Emc is expressed within differentiated neurons. We expressed a membrane-tagged GFP (*UAS:mCD8-GFP*) with the *elav-Gal4* driver to exclusively label differentiated neurons at this stage of development (Robinow and White, 1988; Robinow and White, 1991; Yao and White, 1994). Within the ventral nerve cord (VNC) of third instar larvae we observed extensive Emc protein staining by immunohistochemistry in both neurons (labeled with GFP, arrow in Fig. 1A) and non-neuronal cells (arrowhead in Fig. 1A). These data indicate that Emc is present in both the nucleus and cytoplasm of neurons.

**Figure 1:**
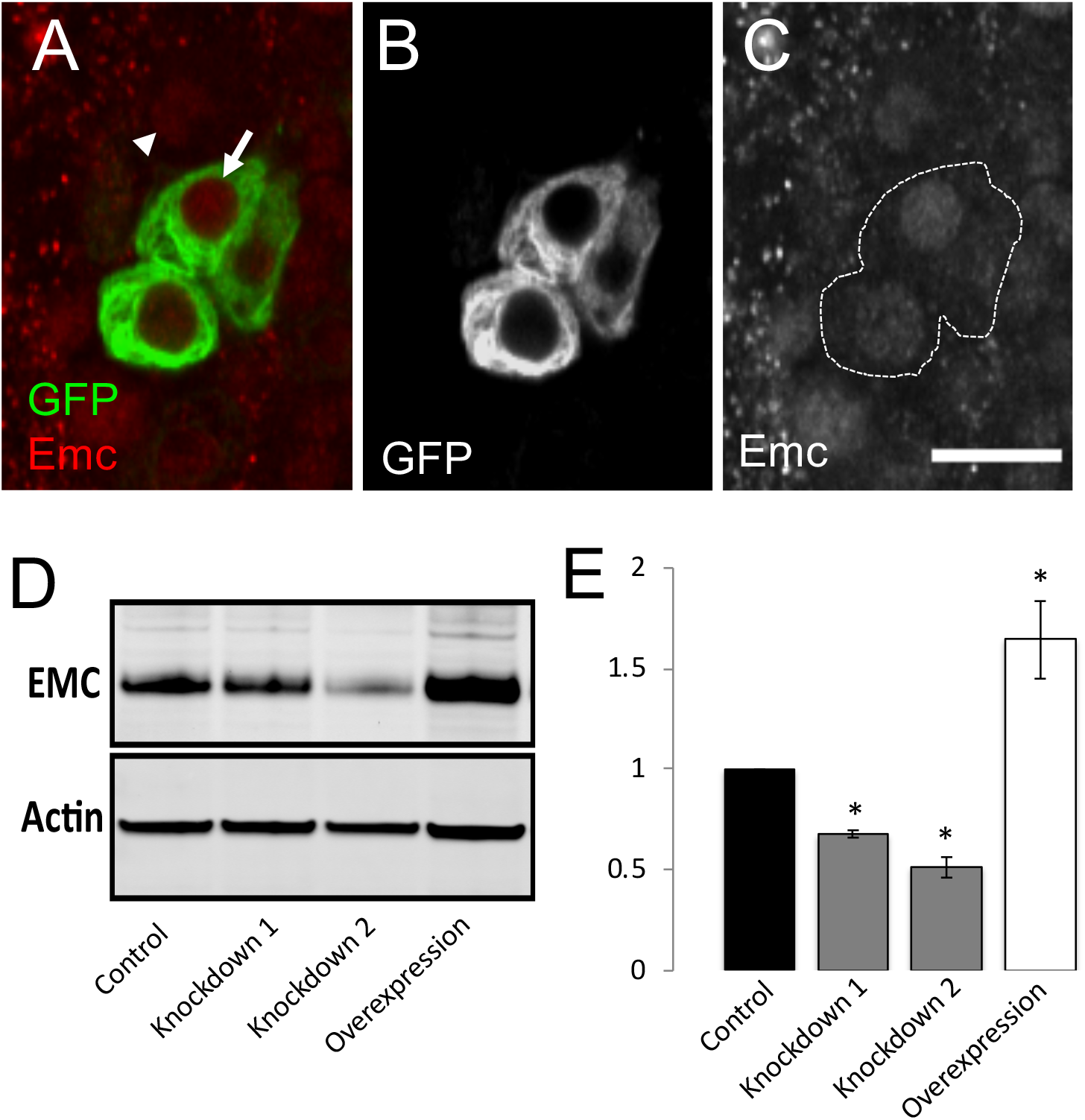
Emc is present in differentiated neurons. Confocal images of third instar larval VNCs labeled with α-Emc (red) and membrane bound GFP (green). Membrane bound GFP labels differentiated neurons. (A-C) WT VNC. (A) VNC labeled for Emc and GFP. Arrow points to neuron (GFP positive). Arrowhead points to non-neuronal cell (GFP negative). (B) GFP channel alone from (A). (C) Emc channel alone from (A). Dotted outline highlights GFP positive cells from (B). Scale bar represents 10 μm. (D) Western blot analysis of Emc protein expression from third instar larval brains. Western blot analysis performed in control, overexpression, and two independent knockdown genotypes. (E) Optical density quantification of western blot analysis compared to actin controls in each genotype. Error bars represent standard error. * *p<0.05*.

We next wanted to determine the function of Emc in neurons. To do this, we knocked down and overexpressed Emc specifically in neurons driving an overexpression reagent (*UAS:emc*) to overexpress Emc and RNA interference (RNAi) to knockdown Emc expression by driving two different short hairpins targeting *emc* under UAS control (*UAS:emcRNAi^1^;* and *UAS:emcRNAi^2^*) with Dicer *elav*-Gal4 (*Dcr2;;elav-Gal4*). In order to validate these reagents, we performed Western blot analysis of Emc protein in third instar larval brains (Fig. 1D). We observed a significant decrease in Emc protein level using both independent RNAi lines targeting Emc (~35% using *UAS:emcRNAi^1^* and ~50% using *UAS:emcRNAi^2^*) compared to wild type controls (Fig. 1D-1E). Additionally, we observed an increase in Emc protein level (~150%) when overexpressed compared to wild type controls (Fig. 1D-1E).

To determine Emc function in neurons, we used the well-defined innervation of the third instar *Drosophila* neuromuscular junction (NMJ), muscle 6/7 of abdominal segment 3, as a model to assess axonal branching and bouton number. We found that Emc overexpression in neurons leads to increased branching and bouton number compared to controls (Figs 2A, 2B, 2E), while knockdown of Emc in neurons leads to decreased branching and boutons compared to controls (Figs. 2A, 2C, 2E). This is the exact opposite phenotype we previously observed with knockdown and overexpression of Da in neurons, respectively (D’Rozario et al., 2016), and is consistent with Emc operating to repress Da function in neurons as it does in mitotic cells. Based on these data, we decided to continue our analysis using *UAS:emcRNAi^2^*.

**Figure 2:**
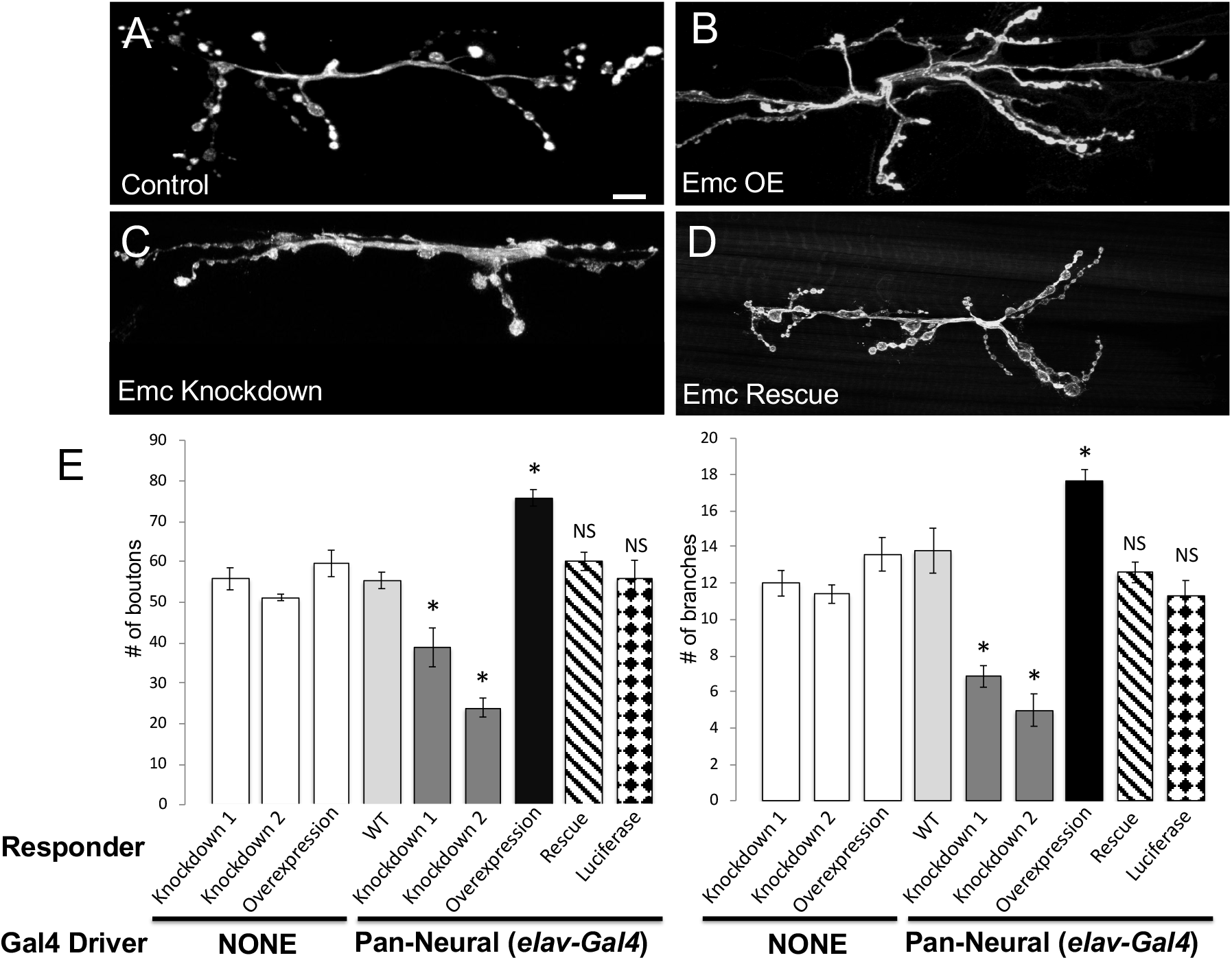
Altered Emc expression affects axonal branch and bouton number. (A-D) Confocal images of third instar larval NMJs, muscles 6 and 7, abdominal segment 3, labeled with α-HRP (white). (A) Control. (B) Emc overexpression genotype. (C) Emc knockdown genotype. (D) Emc rescue (Emc knockdown and Emc overexpression expressed in the same genetic background). (E) Quantification of the total number of boutons and branches. Genotypes indicated. Responder corresponds to *UAS* mediated expression. Gal4 Driver corresponds to *Dcr2;;elav-Gal4* (Pan-Neural). Scale bar represents 10 μm. Error bars represent standard error. * *p<0.05* compared to wild type control. NS is not significant compared to wild type control.

### Emc represses Da function in differentiated neurons

Given the striking NMJ phenotypes observed in larvae expressing Emc knockdown and overexpression, we next wanted to determine the interplay between Emc and Da in the control of branch and bouton number in differentiated neurons. Therefore, we coexpressed four combinations of transgenes using the pan-neural driver to alter Emc and Da expression: 1) Emc knockdown with Da knockdown; 2) Emc overexpression with Da overexpression; 3) Emc knockdown with Da overexpression, and 4) Emc overexpression with Da knockdown. We observed that compared to the decreased branches and boutons in Emc knockdown alone (Fig. 2C), or the increased branches and boutons in Da knockdown alone (Fig. 3C and (D’Rozario et al., 2016)), Emc and Da double knockdown animals displayed an intermediate phenotype (Fig. 3D). While these animals displayed an increased bouton number compared to controls (Fig. 3G), they also displayed significantly fewer boutons than Da knockdown alone (Fig. 3G) and increased boutons compared to Emc knockdown alone (Fig. 2F). Interestingly, these animals did not display significantly altered branch number compared to controls (Fig. 3G), indicating that this manipulation can rescue branching phenotypes associated with knockdown of either Emc or Da alone. Taken together, these data suggest that decreasing Emc and Da together is not as deleterious as decreasing each one individually.

**Figure 3:**
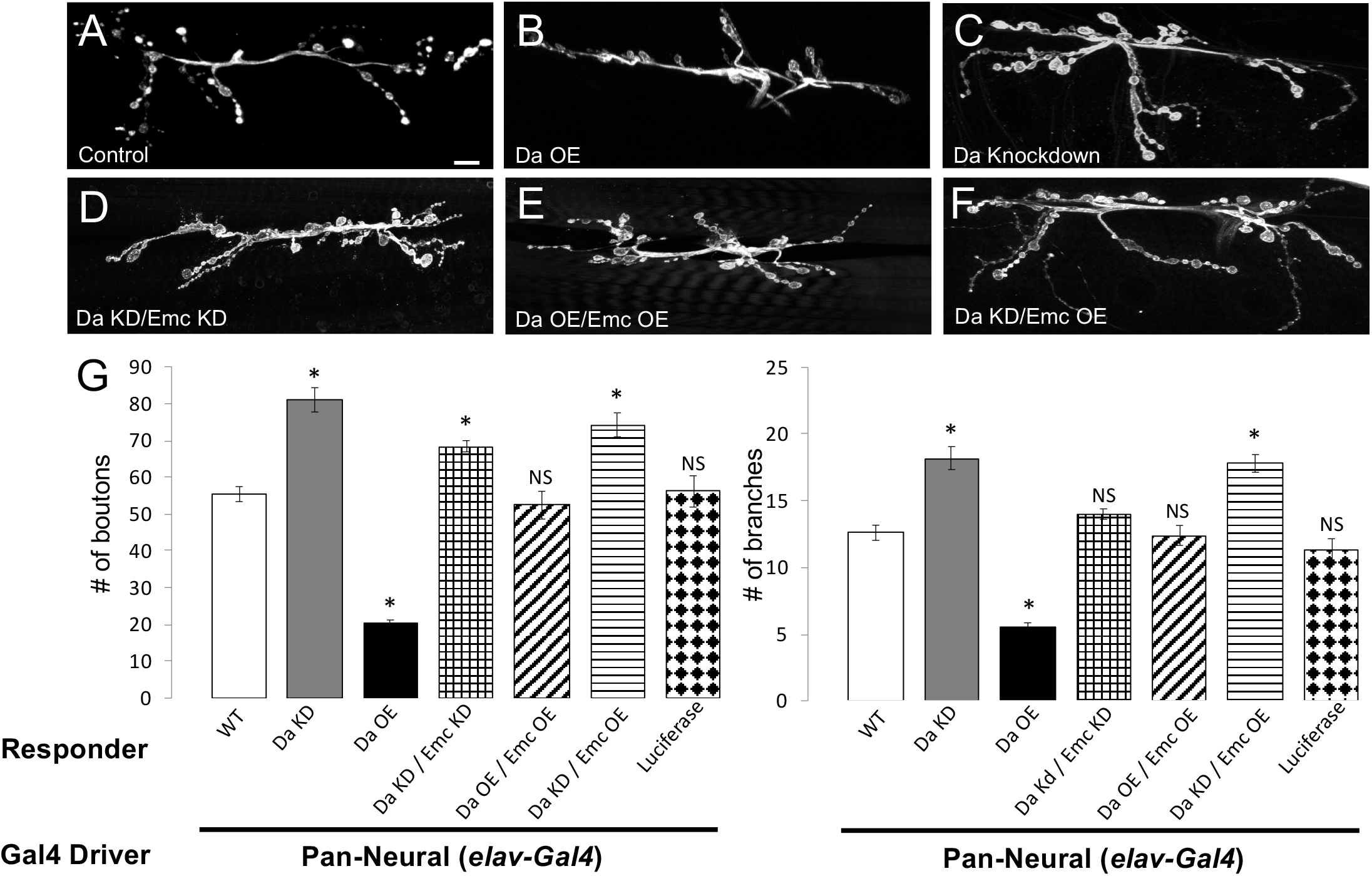
Emc represses Da function in differentiated neurons. (A-F) Confocal images of third instar larval NMJs, muscles 6 and 7, abdominal segment 3, labeled with α-HRP (white). (A) Control. (B) Da overexpression. (C) Da knockdown. (D) Da knockdown and Emc knockdown in the same genetic background. (E) Da overexpression and Emc overexpression in the same genetic background. (F) Da knockdown and Emc overexpression in the same genetic background. (G) Quantification of the total number of boutons and branches. Genotypes indicated. Responder corresponds to *UAS* mediated expression. Gal4 Driver corresponds to *Dcr2;;elav-Gal4* (Pan-Neural). Scale bar represents 10 μm. Error bars represent standard error. * *p<0.05* compared to wild type control. NS is not significant compared to wild type control.

Conversely, we observed that compared to the increased branches and boutons we observed in Emc overexpression alone (Fig. 2B), or the decreased branches and boutons in Da overexpression alone (Fig. 3B and (D’Rozario et al., 2016)), Emc and Da double overexpression animals displayed no significant difference compared to control animals in either the number of boutons or the number of branches (Figs. 3E, 3G). This suggests that increasing Emc and Da together is not as deleterious as increasing each one individually.

Interestingly, Da overexpression coupled with Emc knockdown was lethal. However, Da knockdown coupled with Emc overexpression produced animals that mimicked both an Emc overexpression alone and Da knockdown alone phenotype (Fig. 3F). These animals showed increased branches and boutons compared to controls, which is consistent with Emc overexpression and Da knockdown alone (Fig. 3G and 2F). Taken together, these data suggest that the relative balance between Emc and Da in neurons is important for proper branch and bouton number.

### Altered Emc or Da expression affects motor behavior

To determine if altered Emc protein levels in differentiated neurons has any effect on motor function (the behavior controlled by VNC motor neurons), we analyzed the behavioral output of the transgenic larvae via measurement of larval contraction rate. Larval body wall contraction relies on proper glutamatergic neurotransmission and is significantly affected by mutations that affect transmission (Sandstrom, 2004). Further, we have previously shown that altered Da expression in differentiated neurons leads to synaptic transmission defects (D’Rozario et al., 2016). We observed a significant decrease in contractions when we knocked down or overexpressed Emc in differentiated neurons compared to controls (Fig. 4). We were able to completely rescue this defective behavior when we simultaneously knocked down and overexpressed Emc (Fig. 4). As previously reported, we also observed a significant decrease in motor behavior when we knocked down and overexpressed Da in neurons (Fig. 4) (D’Rozario et al., 2016). However, in each of the Emc/Da double transgenic animals (Emc knockdown/Da knockdown; Emc overexpression/Da overexpression; Emc overexpression/Da knockdown), we observed a significant decrease in motor behavior relative to controls. However, we observed a significant increase in contractions in Emc knockdown/Da knockdown and Emc overexpression/Da overexpression relative to Emc knockdown and Emc overexpression alone, respectively (Fig. 4). Taken together, these data suggest that proper Emc and Da function is required in neurons for proper motor behavior. Further, although alteration of both Da and Emc together showed minimal effects on NMJ bouton and branch number, there is still a significant negative effect on motor behavior when these proteins are altered.

**Figure 4:**
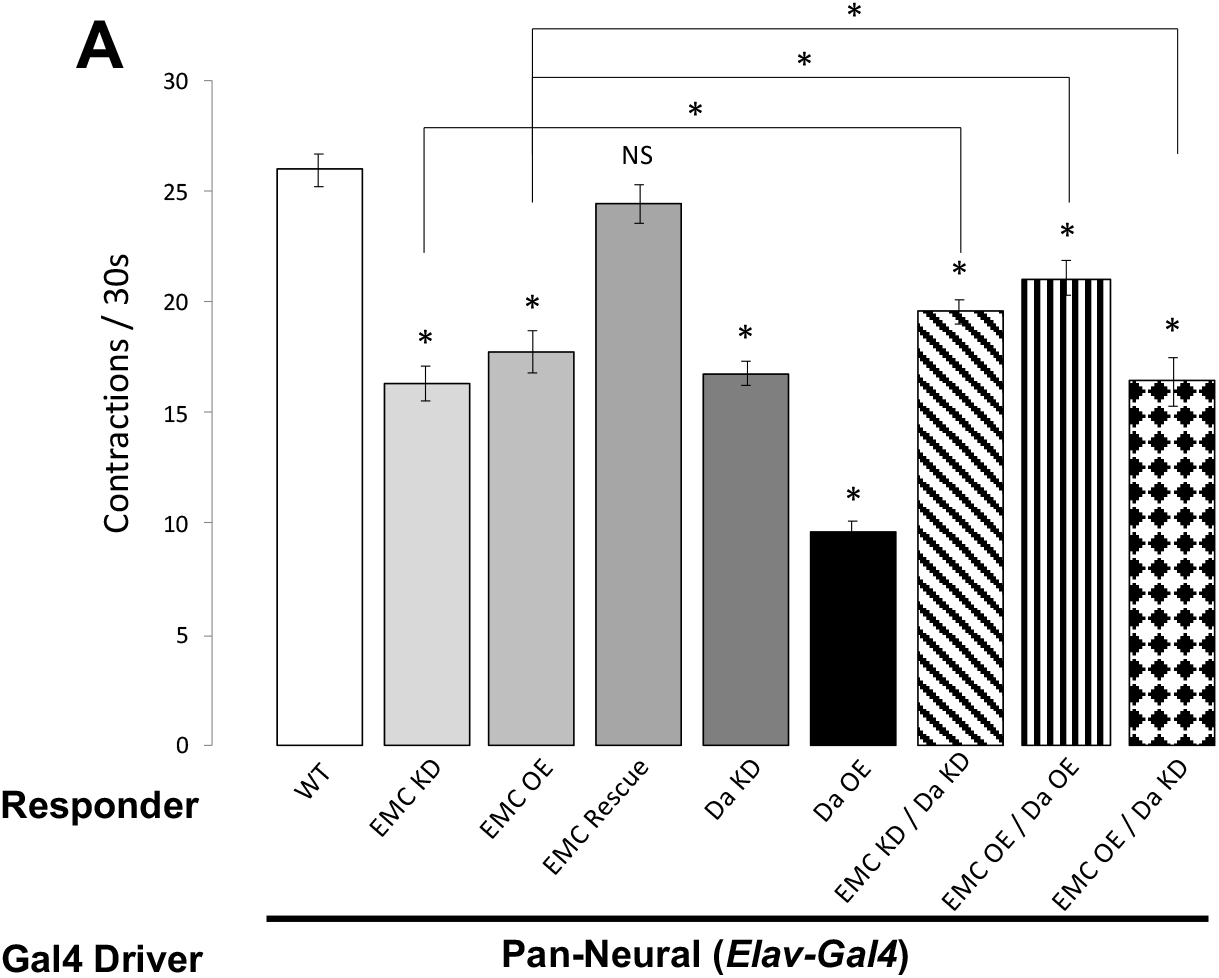
Altered Emc expression affects larval behavior. (A) Total number of larval contractions per 30 seconds in different genotypes. Genotypes indicated. Responder corresponds to *UAS* mediated expression. Gal4 Driver corresponds to *Dcr2;;elav-Gal4* (Pan-Neural). Error bars represent standard error. * *p<0.05*. NS is not significant.

### Emc physically interacts with Da primarily in the cytoplasm of differentiated neurons

Emc belongs to the class V family of HLH proteins which lacks a basic DNA binding domain, but still contains an HLH dimerization domain, and therefore is thought to repress Da function by preventing Da from binding DNA and controlling transcription. Previous work from multiple labs have shown that Da and Emc interact with each other in a number of mitotic cells (Alifragis et al., 1997b; Georgias et al., 1997; Murre et al., 1989b; Spratford and Kumar, 2015b). To assess whether Da and Emc physically interact in differentiated neurons specifically, we utilized the Proximity Ligation Assay (PLA) with established Da and Emc antibodies (Brown et al., 1995; Cronmiller and Cline, 1987; D’Rozario et al., 2016; Spratford and Kumar, 2015a; Spratford and Kumar, 2015b). We expressed each transgene using the *elav-Gal4;UAS:mCD8-GFP* driver to mark differentiated neurons. Surprisingly, we observed a majority of PLA signal in the cytoplasm of these neurons in wildtype conditions (arrows in Figs. 5A-D, 5N) compared to no primary antibody control (Figs. 5E-H). We observed decreased PLA signal upon knockdown of both Emc and Da (Fig. 5M), but no significant effect on PLA signal upon overexpression of either Emc or Da (Fig. 5M).

**Figure 5.**
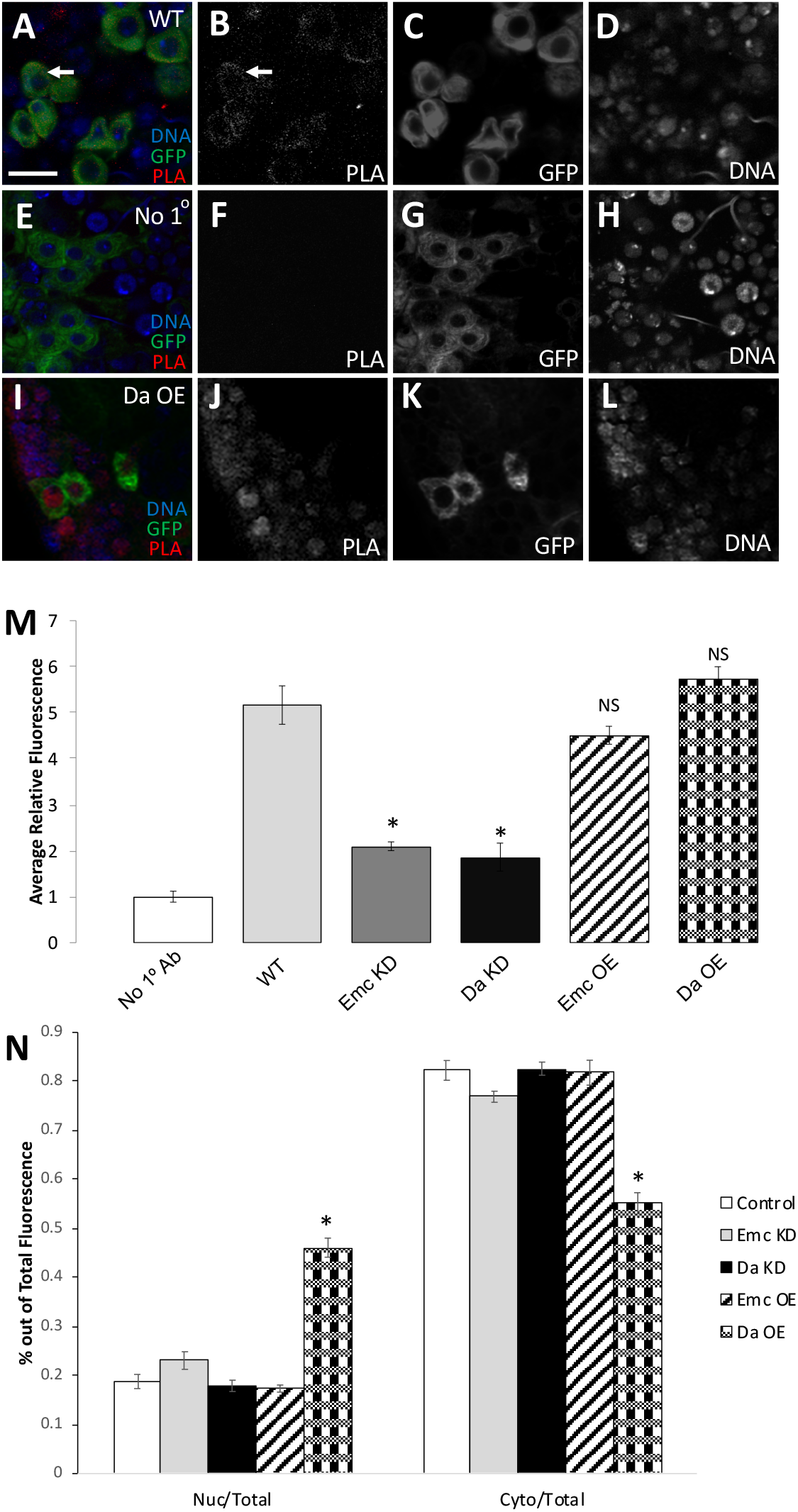
Emc interacts with Da mostly in the cytoplasm. (A-L) 3^rd^ instar VNC neurons labeled with PLA analysis using α-Da and α-Emc (A-D). (E-H) PLA analysis with no primary antibody. (I-L) PLA analysis in Da overexpression background. (B, F, J) are PLA channel alone from (A, E, I). (C, G, K) shows differentiated neurons marked with GFP expression from (A, E, I). Membrane bound GFP labels postmitotic neurons. (D, H, L) shows DNA from (A, E, I). Scale is 10 μm. (M) Shows average relative fluorescence of PLA channel for each genotype indicated. (N) Shows average relative fluorescence of PLA channel in nuclear/total and cytoplasmic/total fluorescence for each genotype listed. Error bars represent standard error. * *p<0.05*. NS is not significant.

We also observed that while a majority of PLA signal was cytoplasmic in a majority of the genotypes we tested (Fig. 5N), a significant amount of PLA signal became nuclear upon overexpression of Da (Figs. 5I-L, 5N). Taken together, these data suggest that Da and Emc also physically interact in neurons, but primarily do so in the cytoplasm of cells in wild type neurons.

### Emc subcellular localization is dependent upon Da expression

Given that we observed an alteration of PLA signal upon Da overexpression, we next wished to determine if the subcellular localization of Emc was dependent upon the expression levels of Da, and vice versa. We quantified the amount of Da present in the nucleus vs. the cytoplasm in wild type neurons and compared this ratio to neurons where Emc was knocked down or overexpressed. We did not observe any significant differences in the amount of Da in the nucleus vs. the cytoplasm compared to controls when Emc was overexpressed (Figs. 6A, 6C, 6G). However, there was a slight, but significant increase in the amount of Da in the nucleus when Emc was knocked down (Figs. 6A-B, 6G) suggesting that total Emc protein may play a role in Da localization to the cytoplasm. Additionally, we did detect a significant increase in the amount of Emc that is present in the nucleus when Da was knocked down compared to controls (Figs. 6D-E, 6H). This altered nuclear/cytoplasmic ratio was even more pronounced when Da was overexpressed in differentiated neurons (Figs. 6F, 6H), consistent with our observation from the PLA analysis. Taken together, these data suggest that Da expression is capable of altering Emc subcellular localization in differentiated neurons.

**Figure 6.**
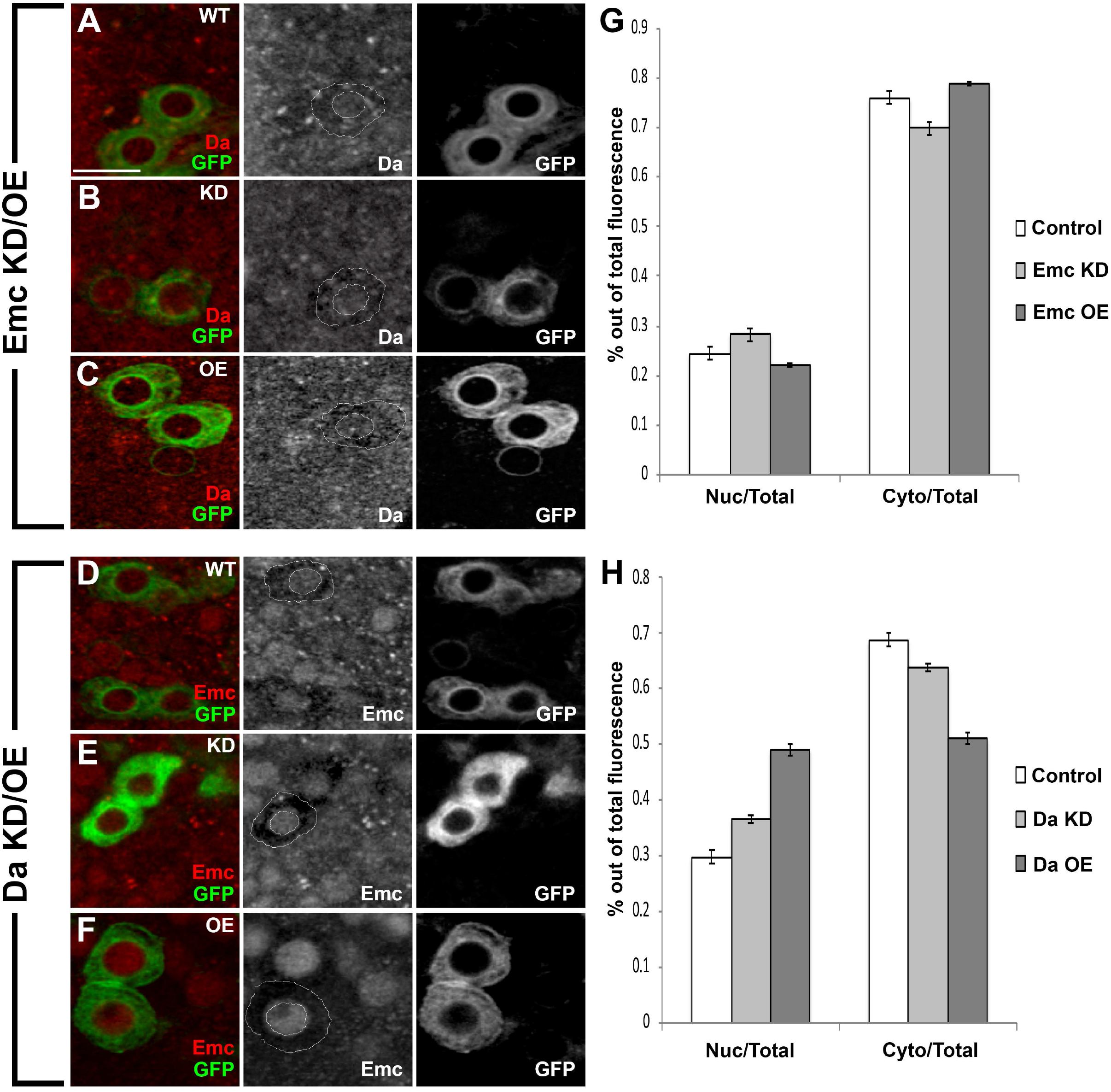
Da expression affects Emc subcellular localization. (A-C) Confocal images of third instar larval VNC labeled with α-Da (red) and membrane bound GFP (green). Membrane bound GFP labels postmitotic neurons. Solid white outline depicts subcellular boundaries of a GFP labeled cell. (A) WT VNC. (B) Emc knockdown VNC. (C) Emc overexpression VNC. (D-F) Confocal images of third instar larval VNCs labeled with α-Emc (red) and membrane bound GFP (green). Membrane bound GFP labels postmitotic neurons. Solid white outline depicts subcellular boundaries of a GFP labeled cell. (D) WT VNC. (E) Da knockdown VNC. (F) Da overexpression VNC. (G) Quantification of the amount of nuclear vs. cytoplasmic fluorescence when Emc is knocked down and overexpressed. (H) Quantification of the amount of nuclear vs. cytoplasmic fluorescence when Da is knocked down and overexpressed. Scale bar represents 10 μm. Error bars represent standard error. * *p<0.05*. NS is not significant.

### Emc promotes *neurexin* expression in differentiated neurons

We had previously shown that Da mediates the repression of NMJ boutons and branches in motor neurons via the repression of the cell surface adhesion molecule Neurexin (D’Rozario et al., 2016). Using yeast 2-hybrid and transcriptional assay studies, Spratford and Kumar have shown that Emc is capable of sequestering Da away from DNA, thus preventing Da-mediated transcription (Spratford and Kumar, 2015b). Our work here suggests that this may be accomplished by Emc sequestering Da in the cytoplasm of neurons. In order to determine if Emc is also capable of controlling *neurexin* expression in neurons via the repression of Da function, we first assessed whether Emc altered *da* mRNA levels. We knocked down and overexpressed Emc specifically in differentiated neurons using *Dcr2;;elav-Gal4*. As a control, we also examined *da* transcript levels in flies expressing the *UAS:Da-RNAi* construct. While we observed a significant decrease in *da mRNA* when we knocked down Da (Fig. 7A), we did not observe any effect on *da mRNA* levels when we either knocked down or overexpressed Emc (Fig. 7A), suggesting that Emc does not alter *da* mRNA levels in differentiated neurons. We next analyzed the level of *neurexin* mRNA. We observed a significant increase in *neurexin* mRNA upon Da knockdown (as we have previously reported (D’Rozario et al., 2016)). We also observed a significant decrease in *neurexin* mRNA levels when we knock down Emc (Fig. 7B), and a significant increase in *neurexin* mRNA levels when we overexpress Emc (Fig. 7B). These data suggest that Emc is required to control the levels of *neurexin* mRNA in differentiated neurons.

**Figure 7.**
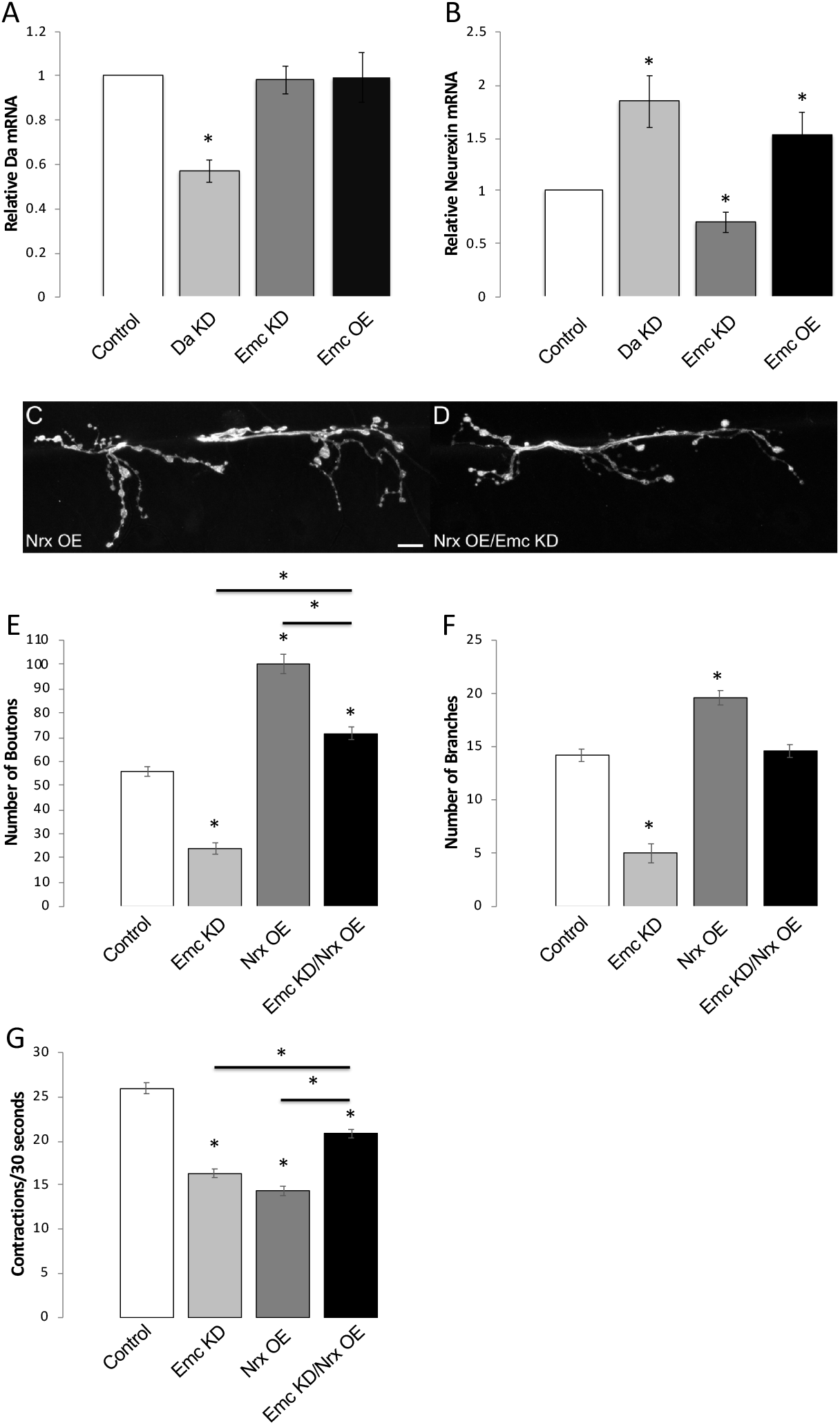
Emc promotes Neurexin expression in neurons. (A) RT-qPCR analysis of *da* mRNA levels from third instar larval brains. RT-qPCR analysis performed in control, Emc overexpression, Emc knockdown, and Da knockdown genotypes. (B) RT-qPCR analysis of *neurexin* mRNA levels from third instar larval brains. RT-qPCR analysis performed in control, Emc overexpression, Emc knockdown, and Da knockdown genotypes. (C-D) Confocal images of third instar larval NMJs, muscles 6 and 7, abdominal segment 3, labeled with α-HRP (white). (C) Neurexin overexpression. (D) Neurexin overexpression and Emc knockdown expressed in the same genetic background. (E-F) Quantification of the total number of (E) boutons and (F) branches. Genotypes indicated. Responder corresponds to *UAS* mediated expression. Gal4 Driver corresponds to *Dcr2;;elav-Gal4* (Pan-Neural). (G) Total number of larval contractions per 30 seconds in different genotypes. Genotypes indicated. Responder corresponds to *UAS* mediated expression. Gal4 Driver corresponds to *Dcr2;;elav-Gal4* (Pan-Neural). Scale bar represents 10 μm. Error bars represent standard error. * *p<0.05*. NS is not significant.

Given that Emc knockdown decreases *neurexin* expression, we wanted to determine if we could effectively rescue the Emc knockdown NMJ and motor behavior phenotypes by overexpressing Neurexin in an Emc knockdown background. Overexpression of Neurexin produced a significant increase in both bouton number and branches compared to the wild type control (Figs. 7C and 7E-7F), consistent with what has been previously shown (Li et al., 2007). However, when we simultaneously knocked down Emc and overexpressed Neurexin, we observed no significant difference in the number of branches between this genotype and the control (Figs. 7D, 7F), and an intermediate phenotype in bouton number compared to Neurexin overexpression alone or Emc knockdown alone (Figs. 7D-E). We next determined whether Neurexin overexpression could also rescue the behavioral phenotypes we observed with decreased Emc. We observed that individually both Emc knockdown and Neurexin overexpression exhibited a significant decrease in motor ability compared to wild type controls (Fig. 7G). However, we observed a significant rescue of this defect when we simultaneously knocked down Emc and overexpressed Neurexin (Fig. 7G). Taken together, these data indicate that Emc also mediates its effect on branch and bouton number by working in opposition of Da to control the levels and function of Neurexin in differentiated neurons.

## DISCUSSION

Class I bHLH proteins require homo- or heterodimerization in order to bind to DNA at E-box consensus sequences and regulate the transcription of target genes (Cabrera and Alonso, 1991; Cronmiller and Cummings, 1993; Massari and Murre, 2000; Murre et al., 1989a; Smith and Cronmiller, 2001). Here, we show that the class V HLH protein Emc promotes axonal arborization and bouton number in neurons by binding Da, and sequestering Da in the cytoplasm of differentiated neurons, thereby preventing Da from binding to and repressing the Da target gene *neurexin*. This work provides novel insights into the molecular functions of class I bHLH and class V HLH proteins in neurons by suggesting that subcellular localization of these proteins may be important in achieving these proteins’ functions.

While we show that Emc sequesters Da away from the DNA by binding to Da predominantly in the cytoplasm, the exact mechanisms of this interaction are unclear. This sequestration may be the result of Emc heterodimerizing with Da in the cytoplasm and preventing Da from entering the nucleus. This explanation is supported by literature showing that the mammalian homolog of Da, Tcf4, is able to enter the nucleus via heterodimerization, even when the Tcf4 protein contains a mutated nuclear localization sequence (Sepp et al., 2011). Thus, blocking the ability of Da to heterodimerize with partners other than Emc may be an important step required to maintain Da in the cytoplasm.

An alternative hypothesis is that Emc is heterodimerizing with Da in the nucleus and exporting Da out of the nucleus. This hypothesis is supported by our data showing that Emc localization increases in the nucleus of neurons when Da is overexpressed. Recently, it was also identified that Da protein levels help regulate Emc protein levels through stabilizing Emc in heterodimers, adding an additional layer of complexity to this interaction (Li and Baker, 2018). Furthermore, Da has been shown to regulate Emc expression via post-translational regulation and transcriptional regulation during adult peripheral neurogenesis (Li and Baker, 2019). Further work will be needed to fully understand the mechanisms underlying the interplay between Da and Emc localization in neurons.

Our work identifies Emc as one binding partner of Da function in differentiated neurons. However, Da is known to have multiple binding partners in mitotic cells and will most likely have additional binding partners that are required for the restriction of axonal branching and bouton number in differentiated neurons as well. Our previous work has shown that Da homodimers also mediate *neurexin* expression in neurons (D’Rozario et al., 2016). However, this does not eliminate the potential for other HLH dimerization partners to be required for normal Da function in neurons. Further, Da has also been shown to affect transcription of target genes through interacting with proteins other than those in than the bHLH family, such as the Notch Intracellular Domain and Suppressor of Hairless in the Notch Transcriptional Complex (Cave et al., 2009). Additionally, in order for Da homodimers to bind to DNA at E-box consensus sequences, additional proteins are required to provide specificity and bind at sites adjacent to E-box sequences (Murre et al., 1994; Tanaka-Matakatsu et al., 2014). Overall, further investigation is required in order to determine the exact proteins required for DNA binding by Da in neurons.

Emc is not the only protein identified that binds to Da and negatively regulates its function. Since Da is heavily regulated post-translationally, other proteins in differentiated neurons could be regulating Da function. Transcription factors such as Nervy, Hairy, Stripe, and Enhancer-of-split proteins have all been shown to bind to Da and repress Da activity (Alifragis et al., 1997a; Fischer and Gessler, 2007; Giagtzoglou et al., 2005; Usui et al., 2008; Wildonger and Mann, 2005; Zhang et al., 2004). Though we have not investigated other potential binding partners here, these proteins (as well as others) may also be involved in controlling Da function in differentiated neurons.

We have shown that Emc controls the levels of *neurexin* mRNA in neurons. However, it is likely that there are more genes whose expression Emc is controlling that regulate bouton and branch number in neurons. Emc could therefore be affecting the expression of additional target genes by regulating Da and/or other neuronal specific bHLH transcription factors. Our previous work identified potential Da target genes in neurons through the analysis of two large-scale genome-wide data sets involving Da and TCF4 (D’Rozario et al., 2016). A list of 44 candidate genes specifically expressed in the *Drosophila* NMJ was generated from this analysis, with *neurexin* being one of the potential targets (D’Rozario et al., 2016). These other 43 genes would make attractive candidates to explore futher to help elucidate the molecular mechanisms underlying how Da restricts axonal arborization and bouton number in neurons.

Emc may also play a role in NMJ branch and bouton number through Da independent functions. For example, Emc has been shown to heterodimerize and negatively regulate class II bHLH proteins during development (Benezra et al., 1990). Additionally, the mammalian homologs of Emc (Id proteins) can promote neurite elongation through heterodimerizing with the bHLH transcription factor NeuroD and repressing its function (Iavarone and Lasorella, 2006).

Haploinsufficiency of *TCF4* has been associated with Pitt-Hopkins syndrome (Amiel et al., 2007; Tamberg et al., 2015), while common small nucleotide polymorphisms in *TCF4* have been associated with schizophrenia (Consortium, 2011). Defects in human ID protein function are associated with a number of disorders in other cell types including eye disease (Fan et al., 2018; Guo et al., 2015), hydronephrosis (Aoki et al., 2004), and bone disease (Fiori et al., 2006). Further investigation into the mechanisms underlying Da function in neurons and how Emc affects Da function could have a profound impact on our understanding of the etiology of these diseases and may provide insights into additional targets for therapeutic avenues.

In summary, we propose that Emc functions in neurons to promote axonal arborization and bouton formation through binding to and inhibiting the function of Da. The interplay between Emc and Da is important for stable and appropriate innervation by neurons via the regulation of *neurexin* expression. Alterations in the normal balance between Emc and Da in neurons leads to defects in neural circuit formation and behavior, as well as altered Emc/Da protein subcellular location. This research identifies the first binding partner of Da in differentiated neurons and opens the door for further research into elucidating the molecular mechanisms underlying Da mediated restriction of axonal branching and bouton number.

## Materials and Methods

### *Drosophila* Stocks and Genetics

*Drosophila* stocks were maintained at 25°C in a 12:12 light:dark cycle with 60% humidity on standard yeast, cornmeal agar medium as previously described (Chakraborty et al., 2011). To drive expression of transgenes in *Drosophila*, the Gal4/UAS bipartite system was used as previously described (Brand and Perrimon, 1993). Stocks were obtained from Bloomington *Drosophila* Stock Center; *UAS:daRNAi* (BL#*26319*), *UAS:emcRNAi^1^* (BL#*42768*), *UAS:emcRNAi^2^* (BL#*26738*), *UAS:LuciferaseRNAi* (BL#*31603*). Overexpression of Da was performed using *UAS:da* (gift from Hugo Bellen as described in (Jafar-Nejad et al., 2006)). Overexpression of Emc was performed using *UAS:emc;UAS:emc* (gift from Justin Kumar as described in (Spratford and Kumar, 2015a; Spratford and Kumar, 2015b)). Overexpression of Neurexin was performed using *UAS:nrx-1* (gift from Swati Banerjee as described in (Banerjee et al., 2017)). All knockdown and overexpression transgenes were expressed using the *Dcr2;;elav-Gal4* (*embryonic lethal abnormal visual system gene trap* (Campos et al., 1987; Robinow and White, 1991; Yao and White, 1994)). For subcellular localization analysis, PLA analysis, and ventral nerve cord imaging, postmitotic neurons were labeled by driving the axonal marker *UAS:mCD8-GFP* with *elav-Gal4*. The Emc rescue genotype was achieved through crossing *UAS:emc;UAS:emcRNAi^2^* with *Dcr2;;elav-Gal4*. Combination genotypes were *UAS:da;UAS:emc, UAS:da:UAS:emcRNAi^2^, UAS:daRNAi/UAS:emcRNAi^2^, and UAS:emc;UAS:daRNAi* and were crossed with *Dcr2;;elav-Gal4*.

### Immunohistochemistry and Antibodies

Larval brains and NMJs were made accessible via larval filet preparations. Wandering 3^rd^ instar larvae of both sexes were obtained and pinned down on Sylgard lined Petri dishes. Larvae were dissected in PBS and were then fixed in 2% paraformaldehyde for 25 minutes. Tissues were then washed in PBS containing 0.1% Triton X-100 (PBT), permeabilized in PBS containing 0.5% Triton X-100 (PBT), and then blocked with 10% normal goat serum (in PBT) for 20 minutes. Primary and secondary antibodies incubations were performed as previously described (Ghosh et al., 2014). Mouse anti-Daughterless (1:40) was a gift from C. Cronmiller (Brown et al., 1996). Rabbit anti-Extramacrochaetae (1:1000) was a gift from K. Jan and Y. Jan (Spratford and Kumar, 2015a). Secondary antibodies (Jackson ImmunoResearch) included goat anti-mouse Cy5 (1:200), goat anti-rabbit Cy5 (1:200), and fluorescein-conjugated HRP (1:100) to label neuronal membranes. Rabbit anti-GFP tagged FITC (1:500) and mouse anti-Gal4 (1:200) were acquired from Thermo Scientific. Dissected larvae were mounted in Vectashield (Vector Labs, H-1000). The 6/7 NMJ of hemi-segments A3 were used for all studies.

Imaging of NMJs and larval brains was performed using an Olympus FluoView FV1000 laser scanning confocal microscope. Image analysis and quantification was performed using ImageJ software. The number of axonal branches, the type of branching, and the number of synaptic boutons, were quantified as previously described (D’Rozario et al., 2016; Mhatre et al., 2014). Tissue fluorescence intensity was calculated as previously described (D’Rozario et al., 2016). GFP intensity was used to normalize tissue fluorescence in all samples.

### Larval Behavioral Assay

Wandering 3^rd^ instar larvae from both sexes were briefly rinsed with PBS and allowed to acclimate for one minute on a 4% agar 100-mm Petri dish. Individual larvae were then transferred to a clean 4% agar plate. All experiments were performed on a clean flat agar surface under constant white light as previously described (D’Rozario et al., 2016). Body wall contractions were counted for 30 seconds for each larva using a Leica Mz 125 stereomicroscope.

### Proximity Ligation Assay

Duolink Proximity Ligation Assay (PLA) kit was obtained from Sigma-Aldrich. Larval brains were made accessible via larval filet preparations. Larvae were dissected in PBS and then fixed in 4% paraformaldehyde for 25 minutes. Tissues were washed in 0.1% PBT for 15 minutes, 0.5% PBT for 15 minutes, and then blocked in Duolink Blocking Solution for 30 minutes at 37°C. Tissue samples was then incubated in primary antibodies overnight at 4°C (mouse anti-Daughterless (1:40) and rabbit anti-Extramacrochaetae (1:1000)). After incubation, samples were washed in Duolink 1x Wash Buffer A twice for 5 minutes at 25 °C. Samples were then incubated in Duolink PLA secondary probes in a 1:5 dilution for 1 hour at 37°C. Samples were then washed in Duolink 1x Wash Buffer A twice for 5 minutes at 25°C. After washing, tissue samples were incubated in Duolink Ligation stock (1:5) containing ligase (1:40) for 30 minutes at 37°C. Samples were then washed in Duolink 1x Wash A Buffer twice for 2 minutes at 25°C. Tissue samples were then incubated in Duolink Amplification stock (1:5) with polymerase (1:80) for 100 minutes at 37°C. Samples were then washed in Duolink 1x Wash Buffer B twice for 10 minutes and then in Duolink 0.01x Wash Buffer B for 1 minute. Tissue samples were mounted in Duolink Mounting Medium. Images were obtained using an Olympus FluoView FV1000 laser scanning confocal microscope. Image analysis and quantification was performed using Adobe Photoshop software. Mean tissue fluorescence intensity was measured, and background fluorescence was subtracted to calculate a mean corrected fluorescence intensity for each sample. GFP intensity was used to normalize tissue fluorescence in all samples.

### Western Blot

Wandering 3^rd^ instar larval brains were collected to prepare lysates. Lysates were prepared from 35 larval brains that were immediately homogenized in RIPA buffer (50 mM Tris, 150 mM NaCl, 1% SDS, 1% NP-40, and 0.5% deoxycholate, pH 8.0) supplemented with protease inhibitor cocktail (EMD Millipore) and 1mM EDTA. The homogenate was centrifuged at 13,000 rpm at 4°C for 30 minutes to collect supernatant for analysis. Samples were stored at −80°C. Samples were prepared using the 4x NuPage LDS sample buffer (Invitrogen, Inc.) containing 0.2% BME (β-Mercaptoethanol, Sigma-Aldrich) and heated to 95 °C for 10 minutes. Equal lysate volumes were loaded into each well of NuPAGE 4–12% Bis Tris Gel (Invitrogen). Gels were run using MES running buffer and transferred to PVDF membrane (Immobilon Millipore) using a semidry transfer apparatus (Owl Scientific) and NuPage transfer buffer (Invitrogen). Membranes were then blocked with 2% BSA PBS blocking buffer for one hour. The membrane was probed overnight at 4°C with blocking buffer containing rabbit anti-Extramacrochaetae antibody and an antibody to β-actin (Sigma-Aldrich). The membrane was then washed four times with 1X PBST for five minutes each. The membrane was probed for one hour at room temperature with goat anti-mouse secondary antibody (IRDye 680LT; LiCor) and goat anti-rabbit secondary antibody (IRDye 800CW; LiCor). Band intensities were quantified using the ImageStudiolite software.

### Quantitative reverse transcription-polymerase chain reaction (RT-qPCR)

50 Wandering third instar larval brains per genotype were dissected in ice-cold phosphate buffer and immediately transferred to RNA Later (Abion) and stored at −80 °C. RNA isolation was performed using phenol:chloroform extraction followed by alcohol precipitation. RNA was stored in DNAse/RNase free water at −80 °C. An adapted version of iTaq Universal SYBR Green One-Step protocol (Bio-Rad) was used (Latcheva et al., 2019) and samples were run on Bio-Rad C1000 Thermal Cycler CFX96 Real-Time system. Primers for *daughterless, neurexin, extramacrochaetae*, and *RP49* mRNA were made using IDT.

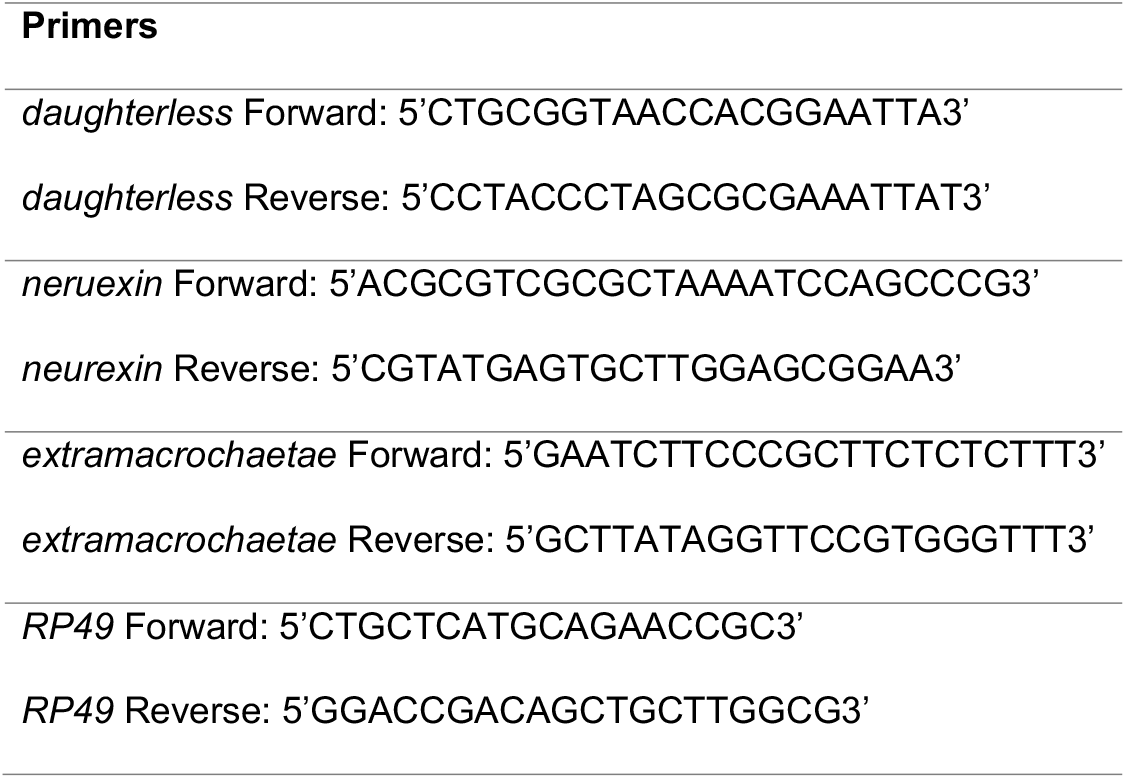

Cycle threshold (Ct) values were obtained graphically for the targets and housekeeping control. ΔCt values were calculated by subtracting the Ct value for each primer set from the Ct value of the house-keeping control. ΔΔCt values were calculated by subtracting the ΔCt value of the control samples from those of the experimental samples. Fold enrichment in expression was as previously described (D’Rozario et al., 2016). Each experiment was performed in triplicate with a minimum of three biological replicates.

## Acknowledgements

We thank Mike Akins, Jennifer Stanford, Felice Elefant, and John Bethea for helpful comments and contributions to this work. We thank Justin Kumar, Swati Banerjee, Hugo Bellen, Mark Biggins, Michael Eisen, the Bloomington *Drosophila* Stock Center, Vienna *Drosophila* Stock Center, and the Developmental Studies Hybridoma Bank for fly stocks, antibodies and reagents; Drexel CIC for imaging analysis and assistance, and Marenda lab members for helpful discussion and comments. This work was sponsored by a grant from the Pitt Hopkins Research Foundation and the Orphan Disease Center at the Perelman School of Medicine at the University of Pennsylvania (to DRM). Work in the Marenda lab is supported by the National Science Foundation IOS 1856439 (to DRM).

